# EGFR-dependent actomyosin patterning coordinates morphogenetic movements between tissues

**DOI:** 10.1101/2023.12.22.573057

**Authors:** D. Nathaniel Clarke, Adam C. Martin

**Affiliations:** Dept.of Biology, Massachusetts Institute of Technology

## Abstract

The movements that give rise to the body’s structure are powered by cell shape changes and rearrangements that are coordinated at supracellular scales. How such cellular coordination arises and integrates different morphogenetic programs is unclear. Using quantitative imaging, we found a complex pattern of adherens junction (AJ) levels in the ectoderm prior to gastrulation onset in *Drosophila*. AJ intensity exhibited a double-sided gradient, with peaks at the dorsal midline and ventral neuroectoderm. We show that this dorsal-ventral AJ pattern is regulated by epidermal growth factor (EGF) signaling and that this signal is required for ectoderm cell movement during mesoderm invagination and axis extension. We identify AJ levels and junctional actomyosin as downstream effectors of EGFR signaling. Overall, our study demonstrates a mechanism of coordination between tissue folding and convergent extension that facilitates embryo-wide gastrulation movements.

## Introduction

Morphogenesis is the development of tissue shape and requires the coordinated movement of thousands of cells^1^. The pattern of coordinated tissue movement is determined by cellular transcription and biochemical signals^2–4^. Regional differences in embryonic cell behaviors can be encoded by the systems that specify the embryonic axes^5^. Because embryos are interconnected three-dimensional structures, shape changes and forces in one region of an embryo often impact other regions^6,7^. Determining how the combination of transcription and signaling differences across an embryo sets up the choreography of different movements across the embryo is pivotal to understanding morphogenesis.

Adherens junctions (AJs) are critical for multicellular behavior, mechanically linking cells and scaffolding actomyosin assembly^8,9^. AJs are composed of conserved complexes of the Cadherin family of adhesion receptors, and the cytoplasmic adaptor proteins β- catenin/Armadillo and α-catenin^10^ (Fig. 1A, B). α-catenin binds filamentous actin (F-actin), thus, the cadherin-catenin complex provides a physical link between the actomyosin cytoskeletons of adjacent cells^10,11^. This linkage is reinforced by additional proteins, such as Vinculin and Afadin/Canoe^12–15^. AJs also serve as platforms for actin and myosin assembly, recruiting both actin nucleators and myosin activators to the zonula adherens^16–19^. In addition to biochemical signals, AJs are regulated by mechanical forces^14,15,20,21^ – tension stabilizes AJs in the *Drosphila* mesoderm^22^. Overall, AJs can be viewed as platforms that integrate signals and are a key control point for regulating tissue-level behavior^9^. The range of mechanisms that control AJs and how they are employed to coordinate tissue movements is an active area of study.

**Figure 1.**
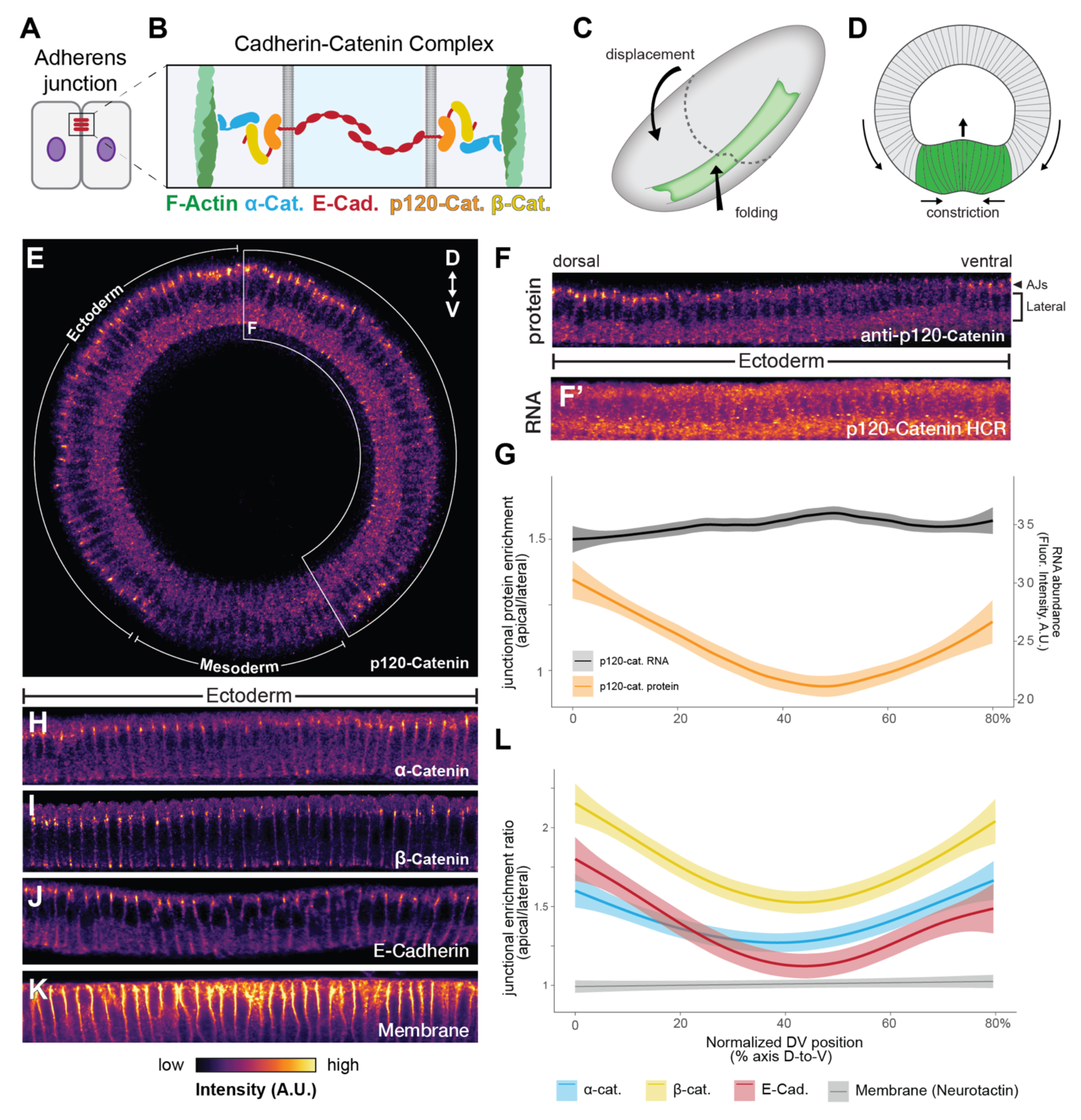
AJ protein levels are patterned along the dorsal-ventral axis. **(A,B)** Schematic of the Adherens junction between two epithelial cells (A), and the Cadherin-Catenin Complex that comprises the functional core of the AJ (B). **(C,D)** Schematic of mesoderm invagination in the *Drosophila* embryo, viewed in whole-mount from a ventrolateral aspect (C) and in cross-section (D). Mesoderm is shown in green, and ectoderm in gray. **(E)** Representative cross-sectional view of a stage 5b embryo stained for p120-Catenin as a marker of the AJ, showing a two-sided gradient pattern from dorsal (top) to ventral (bottom) sides of the embryo. **(F, F’)** Unwrapped projection of p120-Catenin antibody staining from E (F), and a fluorescent in situ hybridization for mRNA of the same gene (F’). **(G)** Quantification of junctional enrichment (ratio of junctional intensity to lateral intensity) of p120-Catenin protein (orange, left axis), and total cytoplasmic p120-catenin mRNA levels (black, right axis). n=5 or more embryos; line represents mean junctional enrichment, plus or minus standard error. **(H-K)** Unwrapped projections of antibody stains for ⍺-Catenin (H), β-Catenin (I), E-Cadherin (J), and Neurotactin (K). **(L)** Quantification of junctional enrichment of AJ factors shown in H-J, in comparison to a general membranal marker. For all quantifications, n=5 or more embryos; line represents mean junctional enrichment, plus or minus standard error.

In animals, including *Drosophila*, separate systems pattern the anterior-posterior (AP) and dorsal-ventral (DV) axes. These axial patterning processes regulate distinct morphogenetic programs. AP axial patterning often regulates convergence and extension movements that extend the anterior-posterior axis of the embryo^23–25^. Convergent extension is driven by planar cell polarized junctional accumulation of non-muscle Myosin II motor protein (myosin), in combination with planar cell polarized protrusions^26–30^. In *Drosophila*, this planar cell polarity of myosin occurs downstream of pair-rule genes, which regulate the expression of Toll-like receptors, Tartan, and the GPCR, Cirl^27,31–34^. DV axial patterning regulates mesoderm internalization and occurs downstream of the ventrally expressed transcription factors Twist and Snail^5,35^. In contrast to axis extension, mesoderm internalization is driven primarily by pulsatile medioapical actomyosin contractility and apical constriction^36–40^. However, planar cell polarity of actomyosin is also observed in mesoderm cell junctions^41^, and pulsatile actomyosin flows are observed in cells undergoing convergent extension^42–45^. Taken together, mesoderm invagination and axis extension are thought to involve largely distinct cellular programs and regulation.

In *Drosophila* mesoderm invagination, local actomyosin contractility constricts the mesoderm and pulls the surrounding ectoderm (Fig. 1C, D). Ultimately the ectoderm closes over the mesoderm. While actomyosin contractility in the mesoderm is thought to be the main driver of its internalization, pushing and stretching by the ectoderm have also been proposed to play important roles^46,47^. The surface cells lost through mesoderm cell internalization have been shown to be compensated for by mitotic rounding and stretching and flattening of dorsal cells^47,48^. In contrast to the dorsal midline cells that flatten, ventral ectoderm cells do not stretch and instead move towards the ventral midline^47^. While differences in cell behavior and adhesion between mesoderm and ectoderm have been described^38,49^, mechanisms regulating the different ectodermal cell behaviors and the interconnection between mesoderm and ectoderm movement are poorly understood.

DV patterning specifies mesoderm and ectoderm and leads to clear differences in AJs between the two cell types^38,49^. Snail expression in ventral mesoderm cells represses E-cadherin transcription and leads to subapical AJs being disassembled^49,50^. DV patterning also partitions different regions of the ectoderm, which is outside of the Snail domain. The intramembrane protease Rhomboid, is expressed in the ventral and dorsal ectoderm, where it activates an EGF ligand^51,52^. Rhomboid expression patterning is critical to the proper cell type specification and development of the *Drosophila* nervous system^53–55^. Downstream EGF signaling is also crucial for specification of cell fates along the dorsal-ventral axis^56,57^. However, EGF’s role in differentiation occurs after gastrulation and whether and how EGF signaling plays a role in gastrulation movements is unknown.

Here, we discovered that AJ levels are patterned in the ectoderm along the DV axis prior to mesoderm invagination. Ectoderm cell AJ levels are regulated by EGF signaling. We show that modulating AJ levels affects junctional actomyosin and that the adhesion/cytoskeletal patterning of the ectoderm plays an important role in the completion of mesoderm internalization. Overall, this work highlights a pivotal role for the regulation of ectodermal cell behaviors in mesoderm invagination and an active role for EGF signaling in morphogenesis prior to its role in nervous system cell fate determination.

## Results

### The Drosophila ectoderm exhibits zones with different AJ levels

To investigate the source of regional differences in ectodermal cell behavior during *Drosophila* gastrulation, we investigated adhesion proteins across the dorsal-ventral (DV) axis. We examined cross sections of fixed embryos stained for a variety of junctional proteins because this enabled antibody staining and high-resolution views of the entire DV axis in single images. Thus, we could compare endogenous protein levels and subcellular localization of AJ proteins across the DV axis. As previously reported, the mesoderm exhibits the lowest levels of junctional proteins (Fig. 1E)^38,49^. In addition, unwrapping the lateral flank of an embryo to display the ectoderm demonstrated a complex pattern of accumulation for all core AJ proteins at the subapical junctions (Fig. 1E – J). The ratio of the subapical to lateral junction intensity for E-cadherin, β-catenin, p120-catenin, and α-catenin all exhibited a double-sided gradient, with the largest peak at the dorsal midline and a second intermediate peak at the ventral ectoderm bordering the mesoderm (neuroectoderm) (Fig. 1G, L). The lateral membrane marker Neurotactin did not display this pattern, with the ratio of subapical to lateral junction intensity being constant across the DV axis (Fig. 1K, L). To determine whether this double-sided gradient of AJ levels represented transcriptional regulation, we performed hybridization chain reaction (HCR) *in situ* hybridization for p120-catenin mRNA (Fig. 1F’), and other AJ genes (Supplemental Fig. 1A). We observed no variation in the level of AJ gene transcripts in the ectoderm of embryos with a clear pattern of junctional protein enrichment, suggesting that AJ protein regionalization was not due to transcriptional control (Fig. 1F’, G). Our results suggest that during gastrulation, the entire AJ complex is modulated across the DV axis in a previously unrecognized pattern within the ectoderm.

### AJ levels are pre-patterned along the DV axis and correlate with EGFR activity

Because the ectoderm stretches in response to mesoderm and endoderm invaginations ^58–62^, it was possible that neuroectoderm cells experience mechanical force that could result in AJ stabilization, as was observed in the mesoderm^22^. Therefore, we hypothesized that the AJ pattern might result from either: 1) mechanical stimulation via the forces of mesoderm cell apical constriction and invagination, or 2) pre-patterning of adhesion from signaling differences along the DV axis. To distinguish between these two hypotheses, we examined the timing of AJ pattern establishment.

We reasoned that if the AJ pattern arose before apical constriction of the mesoderm, it would suggest that increased AJ levels did not result from mechanotransduction, but from pre-patterning. Therefore, we examined embryo cross-sections during cellularization (Fig. 2A), the time when apical-basal polarity and AJs are first established in the blastoderm^63,64^. The timing of each cross section could be inferred from the depth of the invaginating membrane, which has been used to order time points previously^65^. In all regions of the embryo, we observed a basal AJ complex that descended towards the yolk (Fig. 2A - B)^66^. Subapical AJ complexes were assembled as the basal junction complex descended, as previously reported^63^. Subapical AJ assembly initiation was regionalized with AJs in dorsal and ventral ectoderm cells assembling earlier and becoming more intense over time (Fig. 2B - C). Junctional enrichment in ventral neuroectoderm was observed as early as 30 % cellularization (Fig. 2C). These stages occurred well before mesoderm cell apical constriction (Fig. 2A), suggesting that apical constriction forces were not responsible for increasing AJ levels in different regions of the ectoderm.

**Figure 2.**
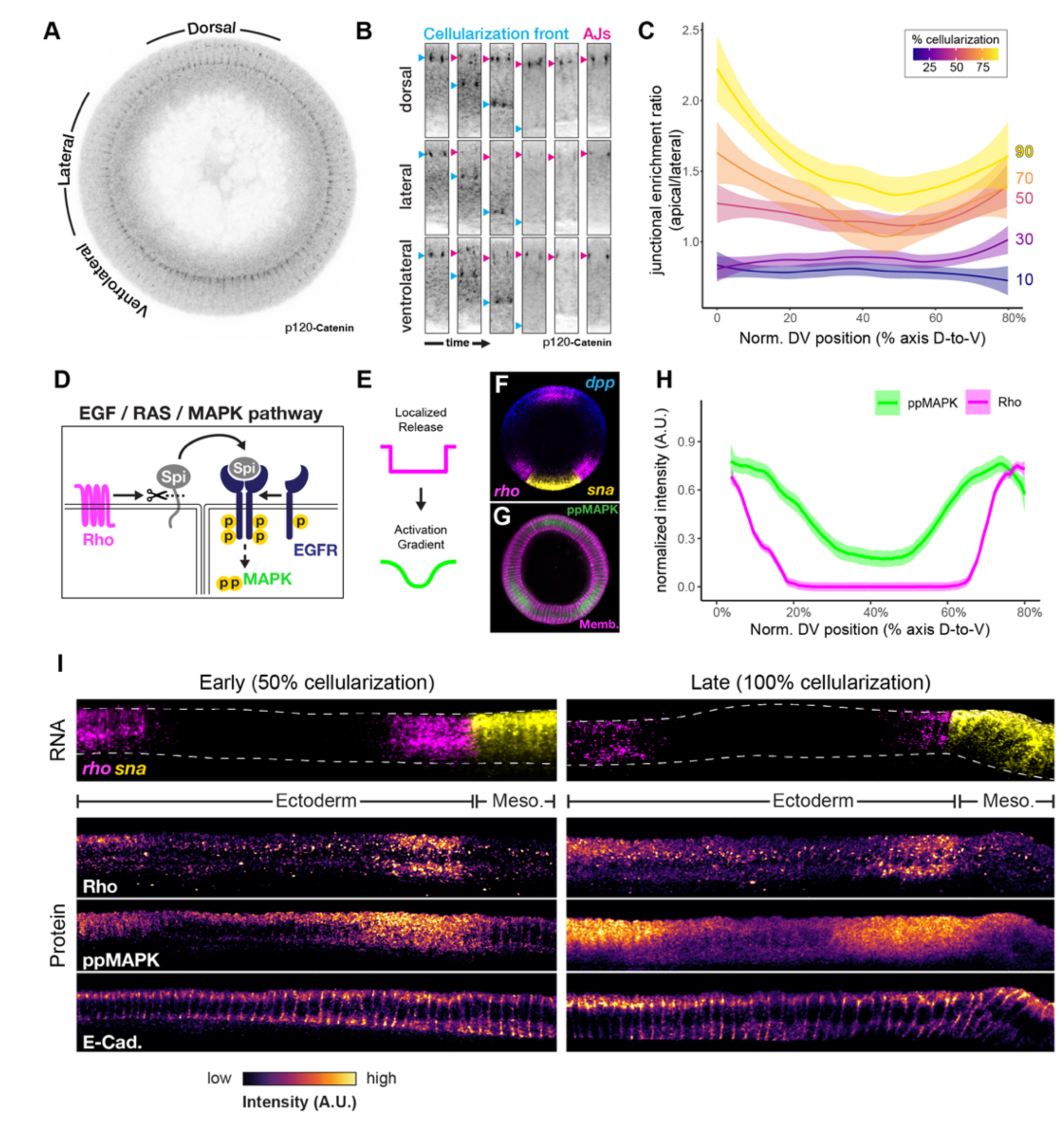
The AJ pattern arises during cellularization and correlates with zygotic EGF signaling activity. (A) Early stage 5 embryo stained for p120-Catenin. Dorsal-ventral regions shown in B are indicated. (B) Representative images of AJ formation in dorsal, lateral, and ventrolateral regions. Basal junctions (cyan arrows) move downwards into the yolk, and AJs (magenta arrows) form in a sub-apical position. (C) Quantification of DV junctional pattern of p120-Catenin over time during junction formation. Colors correspond to percent cellularization, a morphological proxy for time. n=3 or more embryos per stage; line represents mean junctional enrichment, plus or minus standard error. (D) EGF signaling pathway in the *Drosophila* embryo. The integral membrane protease, Rhomboid (magenta), cleaves the membrane-tethered ligand, Spitz (gray), which can act as a short-range morphogen to activate EGFR (blue) and downstream MAPK signaling (green). (E) Schematic of diffusion gradient model for Rho/ppMAPK signaling in the embryo. (F) Fluorescent in situ hybridization for *rhomboid*, the dorsally expressed morphgen *dpp* (BMP), and the mesoderm transcription factor, *snail*. (G) Antibody staining for di-phospho MAPK (green) and membrane (magenta). (H) Quantification of cytoplasmic intensity of *rho* mRNA expression (magenta) in comparison to di-phospho MAPK antibody staining. n=3 embryos; line represents mean junctional enrichment, plus or minus standard error. (I) Unwrapped projections of *rho* mRNA expression patterns in comparison to localization of translated Rho protein, ppMAPK staining, and E-Cadherin, at early (left) and late (right) stages.

To investigate the role of pre-patterning, we noted that the AJ intensity pattern was similar to the published expression pattern of the intramembrane protease Rhomboid (Rho)^67^. Rho cleaves and activates the EGF ligand, Spitz, to promote EGFR signaling (Fig. 2D)^68,69^. We therefore examined EGFR activity using di-phospho Mitogen-activated protein kinase (ppMAPK) staining and compared this to *rhomboid* (*rho*) gene expression and subapical AJ intensity in DV cross-sections. The *rho* gene is expressed in the dorsal and ventral ectoderm, exhibiting sharp expression boundaries (Fig. 2E, F, H). The ppMAPK staining extended beyond these boundaries forming a double-sided gradient, consistent with Rho promoting cleavage and diffusion of EGF ligand (Fig. 2E, G, H). We found a striking spatial correlation between the ppMAPK gradient and AJ levels - both exhibiting a double-sided gradient on the flanks of the ectoderm (Fig. 2I). There was also a temporal correlation between ppMAPK and AJ levels with Rho protein, ppMAPK, and AJ levels, with all becoming higher in ventral neuroectoderm prior to dorsal midline enrichment (Fig. 2I). *EGFR* expression at gastrulation stage was also patterned, being restricted to the ectoderm and exhibiting varying intensity along AP, similar to pair-rule gene stripes, suggesting that there is zygotic EGFR expression at this time (Supplemental Fig. 1B-C). *rho* expression also exhibited AP stripes that were restricted in DV (Supplemental Fig. 1B). Furthermore, EGFR protein colocalized with E-cadherin at AJs (Supplemental Fig. 1D). Taken together, *rho* and *EGFR* expression and ppMAPK activity patterns mirror the emergence of the AJ pattern prior to apical constriction, suggesting a pre-patterned DV AJ intensity gradient.

### DV axial patterning and EGFR signaling are required for AJ pattern formation

To test whether the pattern of AJ intensity depends on DV patterning, we examined AJ levels in cross-sections of embryos dorsalized to varying extents. To do this, we targeted the Toll receptor by maternal RNA interference (RNAi, Supplemental Fig. 2A). RNAi at the standard temperature (25°C), resulted in embryos that uniformly expressed the lateral gene *short gastrulation* (*sog*) around the embryo circumference, suggesting a uniform lateralized cell fate (Supplemental Fig. 2B-D). In contrast, RNAi at elevated temperatures (i.e., 29°C) leads to a more severe knock-down, as reported previously^70^. Toll depletion at higher temperatures resulted in dorsal genes (i.e., *egr*) being expressed around the entire circumference and uniformly high ppMAPK and other dorsal markers, suggesting full dorsalization (Supplemental Fig. 2B, C, E, F, F’’, G, G’’). We found that fully dorsalized embryos exhibit high levels of AJs and ppMAPK without a gradient (Supplemental Fig. 2F - H). In contrast, lateralized embryos exhibited low ppMAPK and lower AJs with uniform intensity around the circumference (Supplemental Fig. 2G’, H’). Overall, these results demonstrate that AJs levels are under control of the DV patterning system and scale with ppMAPK levels.

Because EGF signaling is under control of DV patterning and AJ intensity in wild-type, dorsalized, and lateralized embryos is correlated with ppMAPK levels, we next tested whether EGFR was required to modulate ectodermal AJ levels. We first tested the effects of EGFR depletion. Using maternal and zygotic RNAi and a zygotic EGFR mutant, we found that EGFR depletion results in reduced AJ levels (Fig. 3A – C, Supplemental Fig. 3A). Although the enrichment in the ventral neuroectoderm was eliminated, we continued to observe a one-sided gradient around the dorsal peak (Fig. 3D). We obtained similar results depleting EGFR activity by maternally overexpressing the EGFR inhibitor Argos, or a dominant-negative version of EGFR, with GAL4/UAS (Supplemental Fig. 3B - D). Overall, these results suggested that EGFR is the primary input regulating AJs in the ventral ectoderm region, but that additional inputs regulate dorsal AJs.

**Figure 3.**
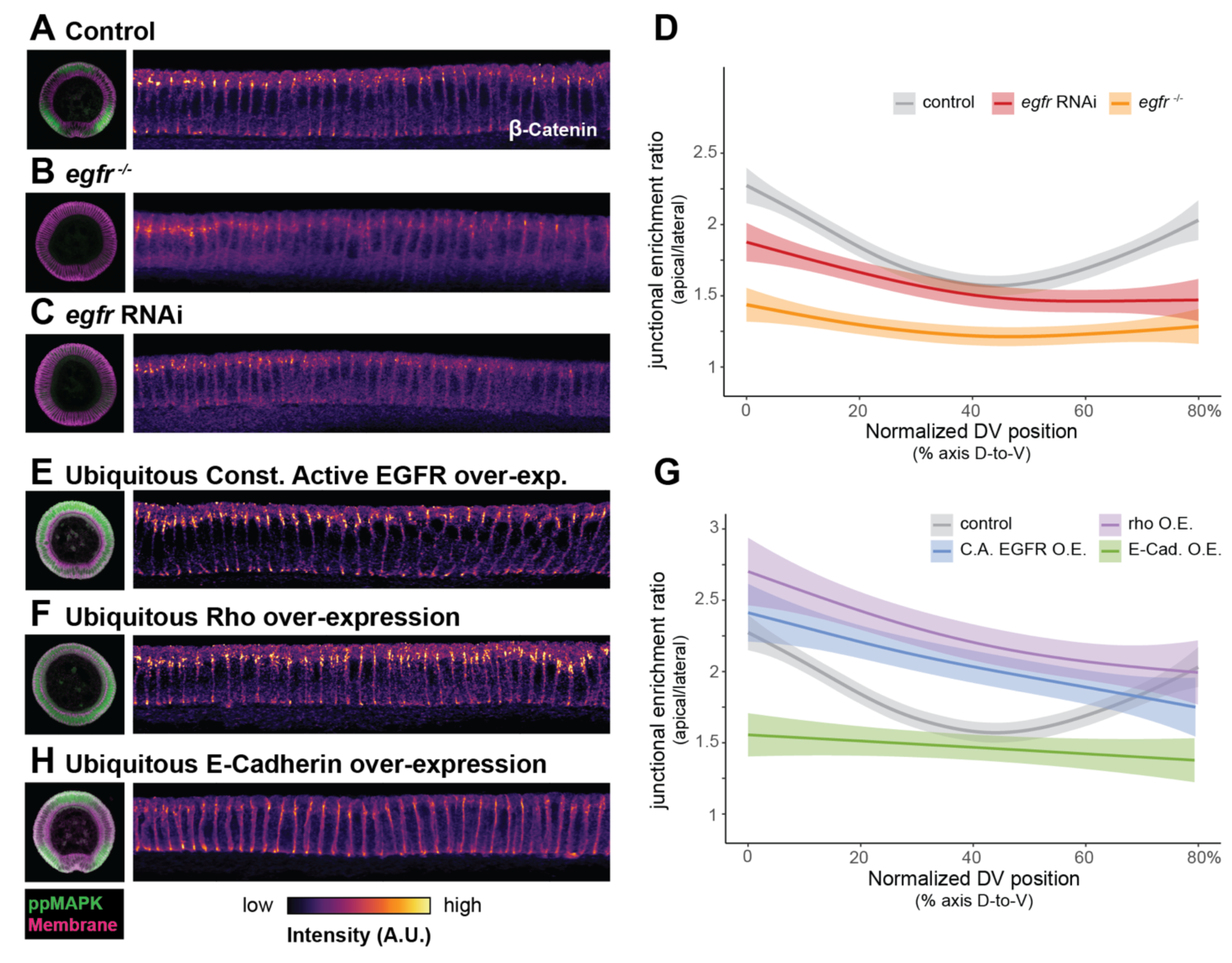
– Disruption of EGF signaling modulates the AJ protein gradient. (A-C) Representative unwrapped projections of anti-β-Catenin staining in EGFR zygotic null mutant (B), and EGFR RNAi knockdown (C) in comparison to control, (A). Cross-sectional view of anti-ppMAPK staining is shown at left to show EGF activation state. (D) Quantification of conditions shown in A-C. For all quantifications, n=5 or more embryos; line represents mean junctional enrichment, plus or minus standard error. (E-F) Representative unwrapped projections of anti-β-Catenin staining under EGF-activating conditions: over-expression of constitutively active EGFR (E), and ubiquitous expression of Rho (F) enrich AJ proteins at sub-apical junctions. (G) Quantification of conditions shown in E-F, and H. For all quantifications, n=5 or more embryos; line represents mean junctional enrichment, plus or minus standard error. (H) Representative unwrapped projection of anti-β-Catenin staining in embryos with ubiquitous over-expression of E- Cadherin.

To perform the converse experiment and activate EGFR, we ubiquitously overexpressed either Rho or a constitutively active form of EGFR (Supplemental Fig. 3A). Ubiquitous EGFR activation led to elevated ppMAPK throughout the DV axis of the embryo (Fig. 3A, E, F). Strikingly, we found that this led to elevated AJ levels through the entire ectoderm, although there was still a one-sided gradient of AJs with a dorsal peak (Fig. 3E - G). Similarly, alternative methods to activate EGFR or downstream Ras signaling resulted in elevated AJs, although activated Ras did not fully mimic the effects of activated EGFR, suggesting that a component of the EGFR signal to AJs occurs in parallel to Ras (Supplemental Fig. 3B, E, and F). In comparison, ubiquitous over- expression of E-cadherin resulted in spreading of AJ proteins across the lateral membrane, resulting in a lower junctional enrichment ratio (3G, H). Taken together, our results suggested that EGF activity leads to elevated assembly and/or stability of the AJs, which leads to higher net AJ levels in ectoderm cells. Our data argue that multiple inputs control AJs in the dorsal ectoderm, but that EGF is the primary input controlling AJ enrichment in the ventral neuroectoderm.

### EGFR signaling is required for mesoderm internalization and germband extension

To determine how EGFR-dependent modulation of adhesion affects tissue flow in the ventral ectoderm (the neuroectoderm), we analyzed cell movement along AP and DV axes. First, we analyzed germband extension along the AP axis (Fig. 4A). Germband extension has been separated into two phases, an initial fast phase (Phase 1, or GBE 1) with rapid extension over ∼ 40 min and a later slow phase (Phase 2) continuing for 20-70 minutes after the rapid extension (Supplemental Fig. 4A)^34,71^. Analysis of EGFR-depleted embryos indicated a slower rate of phase 1 germband extension while phase 2 was unaffected (Supplemental Fig. 4B, C). In contrast to depletion, constitutive activation of EGFR resulted in more rapid elongation during phase 1 of germband extension (Supplemental Fig. 4B, C). Thus, our data suggest that EGFR activity tunes axis extension rate in *Drosophila* before its known role in central nervous system patterning.

**Figure 4.**
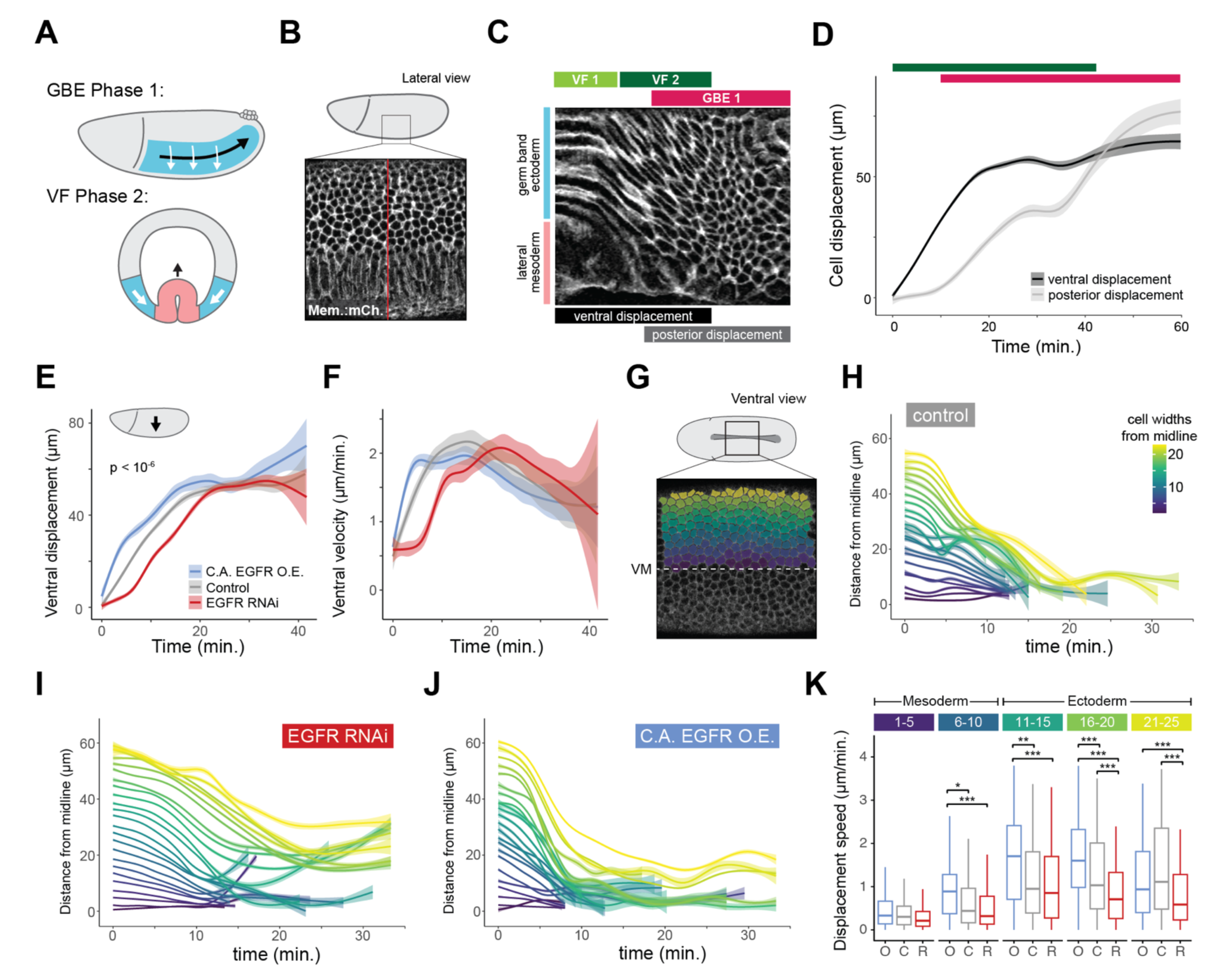
– Disruption of EGFR inhibits cell flow during gastrulation. (A) Schematic of temporally overlapping movements of Phase 1 of germ band extension (GBE) and Phase 2 of ventral furrow formation (VF). Ectoderm and mesoderm are shown in blue and pink, respectively. Arrows indicate approximate direction and speed of cell movement. (B) Schematic of lateral imaging strategy for confocal microscopy. Live imaging marker is ubiquitously expressed Gap43:mCherry. Red line indicates the line scan used to generate the kymograph in C. (C) Annotated kymograph of the GBE process, viewed from a ventrolateral aspect. Tissue regions (left), gastrulation phases (top), and cell movement patterns (bottom) are indicated. Time moves from left to right. (D) Quantification of ventral (black) and posterior (gray) cell displacements measured from cell tracking data of wildtype embryos. Bars at top indicate gastrulation phases. N=3 embryos. Line represents mean plus or minus standard deviation. (E-F) Quantification of ventral displacement (E) and velocity (F) over time in ventrolateral aspect movies of EGFR perturbation conditions (colors same as B). Significance calculated by Kruskal-Wallace test. N=3 embryos each. Lines represents mean plus or minus standard deviation. (G) Schematic of ventral imaging strategy for confocal microscopy. Color gradient overlay shows example cell segmentation and row-wise cell identification strategy used for quantifications. Live imaging marker is Gap43:mCherry. The ventral midline (VM) is indicated. (H-J) Row-wise quantification of cell displacement towards the ventral midline over time in control embryos (H) in comparison to EGFR RNAi knockdown (I) or constitutive EGFR over-expression (J). Colors correspond to cell row, with the mesoderm-ectoderm boundary in green. N = 3 embryos per condition. Lines represents mean plus or minus standard error. (K) Quantification of cell velocity towards the midline in binned cell rows. O, C, and R refer to EGFR over-expression, control, and EGFR RNAi, respectively.

Mesoderm invagination or ventral furrow formation (VF) has also been separated into two distinct phases: the first phase (VF 1) includes apical constriction and furrow formation and the second phase (VF 2) involves the rapid internalization of mesoderm cells and ventral-directed movement of ectoderm cells that close over mesoderm (Fig. 4A – C). The second phase of mesoderm invagination temporally overlaps the first phase of germband extension (Fig. 4A – D). To determine the possible effect of EGFR on mesoderm invagination, we measured the ventral-directed movement of ectoderm cells during phase 2 of mesoderm invagination. Inhibition of EGFR prior to gastrulation reduced neuroectoderm movement towards the ventral midline (Fig. 4E, F), delaying closure of the mesoderm from the surface (Supplemental video). Importantly, inhibition of EGFR did not affect mesoderm cell apical constriction or apical myosin accumulation, consistent with the lack of EGFR expression and downstream activity in the mesoderm (Supplemental video, see below). In contrast to inhibition, EGFR activation accelerated ventrally directed flow. To determine whether specific cell populations in the ectoderm were affected we measured cell displacement as a function of distance from the ventral midline, by cell row (Fig. 4G-K). The mesoderm half-width is 8-10 cells, thus we could distinguish between mesoderm cell movement resulting from apical constriction from the ventrally directed flow of the adjacent ectoderm (>10 cells from midline). We found that EGFR depletion significantly reduced the ventral movement of cells 16 – 25 cell rows from the ventral midline, demonstrating that apical constriction occurs normally and that EGFR inhibition specifically impeded ectoderm movement towards the ventral midline (Fig. 4I, K). In contrast, EGFR activation accelerated the ventral movement of cells from 5-20 cell rows from the ventral midline (Fig. 4J, K). EGFR activation possibly accelerated movement within mesoderm by reducing resistance to apical constriction (see below). Taken together, our results demonstrate that ectodermal EGFR activity modulates the ectoderm ventral movement during late stages of mesoderm invagination and early germband extension.

### EGFR signaling regulates ectodermal cell shape during mesoderm invagination

We hypothesized that patterns of AJ levels in the ectoderm could affect cell mechanical properties in a manner that would explain the observed differences in cell movements. A change in cell mechanics would be expected to alter the balance between cell movement and shape change that accompanies pulling forces by mesoderm cell invagination. To test whether EGFR-mediated regulation of adhesion was important for ectoderm cell shape during mesoderm invagination, we quantified cell aspect ratio at different positions from the mesoderm-ectoderm boundary. We fixed embryos and stained for *snail* and *sim* using HCR *in situ* hybridization in combination with a membrane label (Fig. 5A – D). Marking cell position relative to the single cell row of *sim* expression (row 0), we found that the *egfr* mutant embryos resulted in cells that were more anisotropically stretched along the DV axis (Fig. 5E – F). We also saw a greater incidence of Snail-expressing cells that were left on the embryo surface after mesoderm invagination in the *egfr-/-* mutant embryos, further supporting the role for ectodermal EGF signaling in mesoderm invagination (Fig. 5G). To confirm that EGF signaling regulates ectodermal cell shape, we performed live imaging of the ectoderm during mesoderm invagination with a membrane marker. We measured the average cell area strain in the neuroectoderm and found that there was greater strain in EGFR depleted embryos than in controls (Fig. 5H – I). Taken together, our results demonstrate that EGF signaling changes ectodermal cell shape and movement during mesoderm invagination, which is required to complete mesoderm internalization.

**Figure 5.**
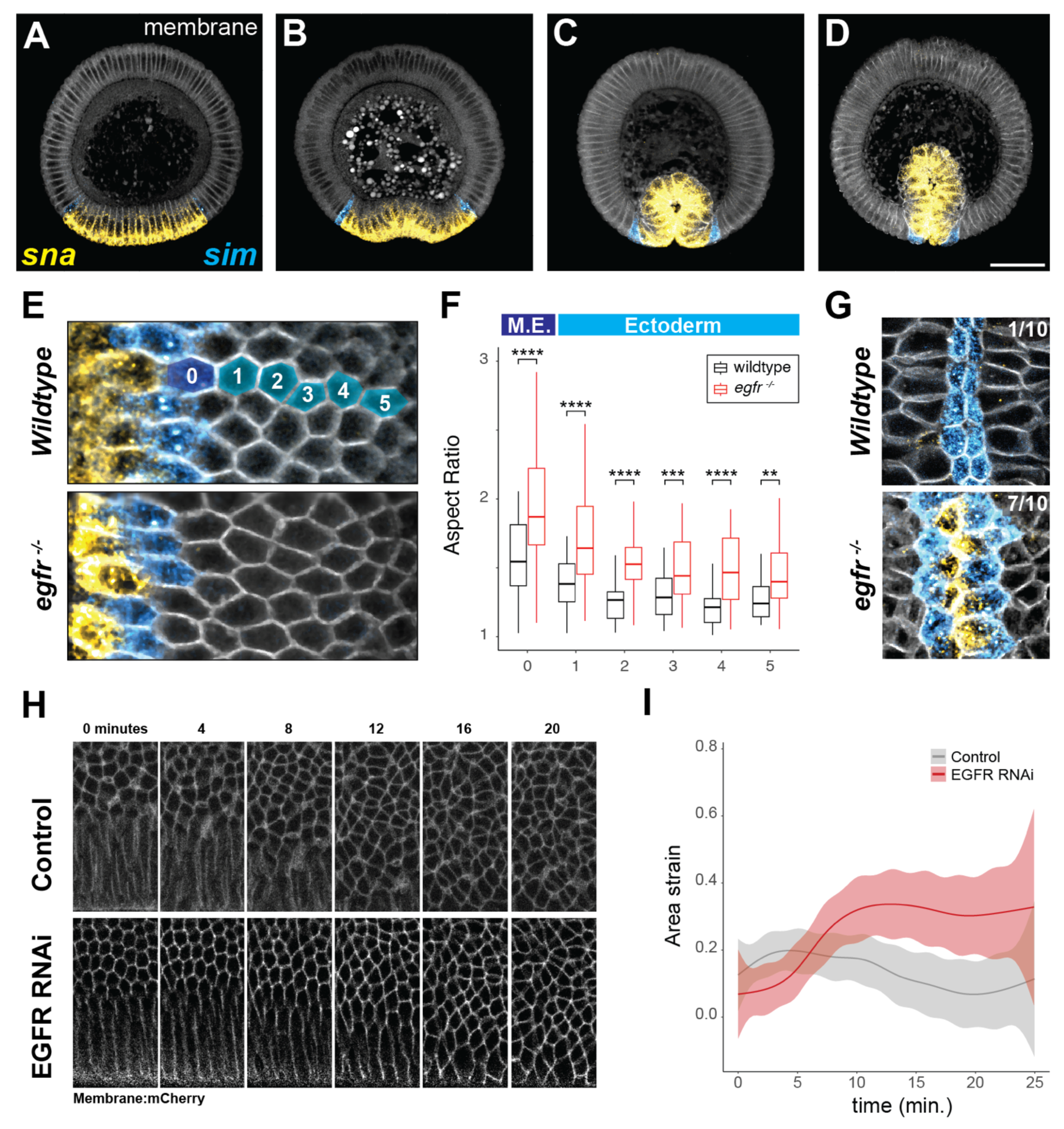
– Modulation of the AJ gradient alters cellular strain in cells adjacent to the folding mesoderm. (A-D) Cross-section series of fluorescent *in situ* hybridizations spanning the ventral furrow formation process. *snail* (*sna*, yellow) marks the mesoderm, and *single-minded* (*sim*, cyan) marks the mesectoderm boundary between mesoderm and ectoderm. Embryos are co-stained with anti-neurotactin antibody to show cell membranes. (E) Comparison of apical views of mid-stage 6 embryos (equivalent stage to C) of wildtype, or zygotic null *egfr* mutant genotypes. Staining is the same as in A-D, and the cell rows used for analysis in F are indicated. (F) Quantification of apical area aspect ratio versus distance from the mesectoderm for wildtype and *egfr* null embryos. (G) egfr zygotic null mutant embryos fail to fully internalize mesoderm. Embryo preparation is the same as in E, but at a later, post-folding stage. Count in upper right indicates number of embryos with *sna*-positive cells on the surface post-folding. (H) Representative live-cell timelapse in control and EGFR RNAi embryos, showing delayed displacement and increased apical area deformation in EGFR RNAi. (I) Quantification of area strain over time from live-cell tracking data. N = 3 embryos per condition. Lines represent mean plus or minus standard deviation.

### EGFR-dependent AJ elevation promotes increased ectodermal actomyosin levels

EGF signaling could impact ectodermal cell shape through direct effects on cell adhesion, which can influence tissue fluidity^72,73^. Alternatively, it was possible that AJs impact cell shape through their scaffolding function and could be modulating junctional actomyosin levels. To address this possibility, we examined F-actin and myosin levels along the DV axis. We found that both the junctional enrichment ratio of F-actin and myosin levels exhibited a double-sided gradient pattern along the DV axis, similar to that of ppMAPK and AJs (Fig. 6A - C). The myosin junctional enrichment ratio does not necessarily capture the total overall myosin density and the ventral-dorsal myosin gradient observed using light sheet microscopy^47,60^. To determine if this actomyosin pattern also depended on EGF signaling, we examined myosin intensity in EGFR depleted and constitutively activated conditions. We found that EGFR depletion disrupted junctional myosin levels, specifically in the neuroectoderm (Fig. 6D, E). In contrast, ubiquitous EGFR activation increased junctional myosin levels throughout the embryo (Fig. 6D, E). *En face* views of junctional actomyosin demonstrated that modulation of the EGF pathway affected the levels of junctional myosin that was recruited to DV cables – with activation of EGFR maintaining myosin planar cell polarity (Fig. 6F). Quantification showed that disrupting EGFR signaling affected junctional myosin, but not medioapical myosin (Fig. 6G). These results demonstrated that EGFR coordinates the regulation of AJs and junctional actomyosin across the DV axis, with additional signaling inputs regulating the activity of these systems at the dorsal midline.

**Figure 6.**
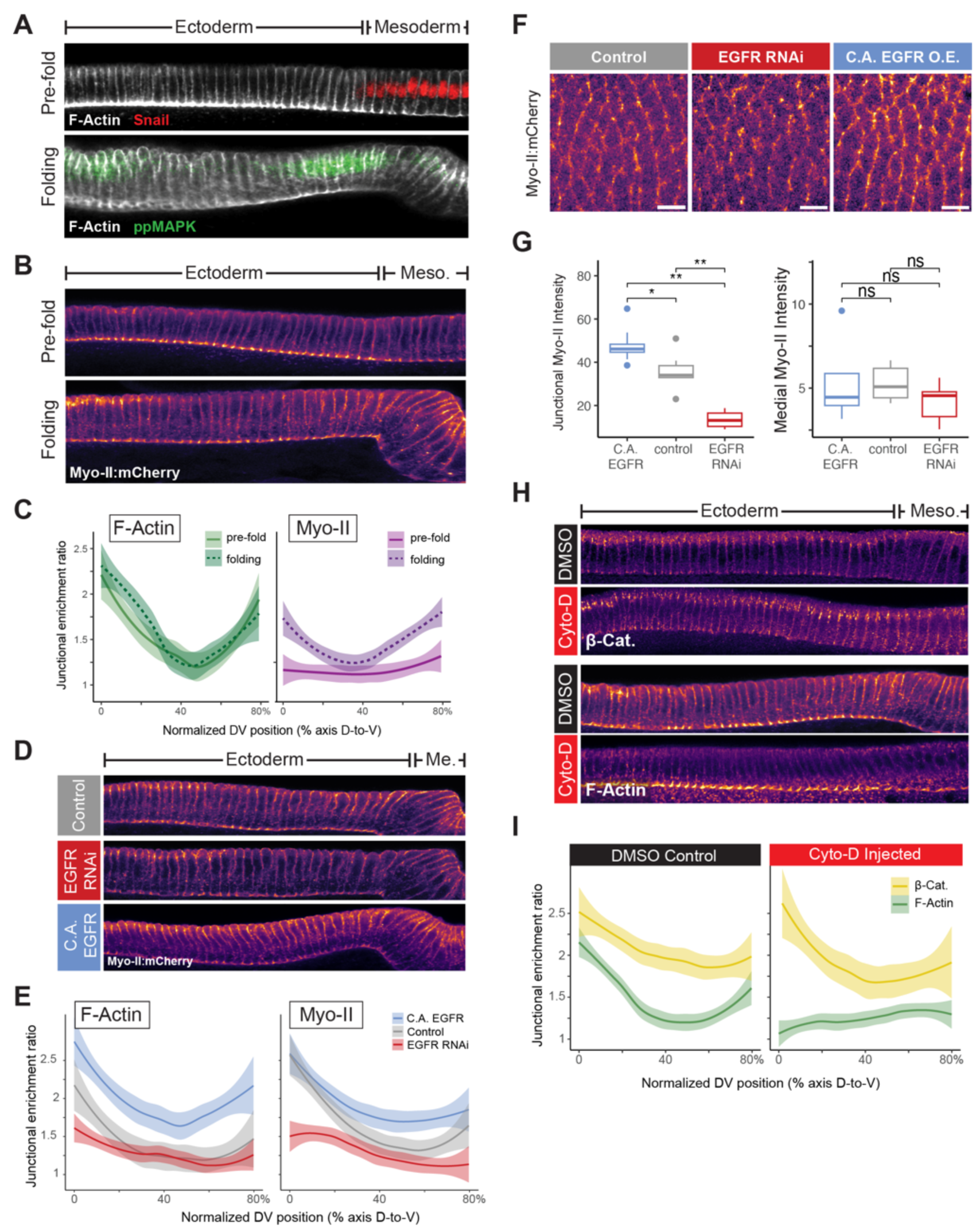
– AJ protein levels modulate actomyosin recruitment in the ectoderm. (A – B) Representative unwrapped projections showing patterns of F-Actin (A), and Myosin (B) distribution during pre-gastrulation (pre-fold) and mid-gastrulation (folding) timepoints. Snail (red) and ppMAPK (green) are shown in A to indicate the mesoderm, and zones of EGF activity, respectively. (C) Quantification of junctional F-Actin and Myosin intensity at pre- and mid-gastrulation timepoints, as shown in A. N=5 embryos. Line represents mean plus or minus standard error. (D) Representative unwrapped projections of endogenous Myo-II:mCherry fluorescence in embryos expressing constitutively active EGFR or EGFR RNAi, in comparison to control. (E) Quantification of junctional F-Actin and Myosin in the EGFR perturbations shown in D. N=5 embryos. Line represents mean plus or minus standard error. (F) en face views of endogenous Myo-II:mCherry fluorescence in embryos expressing constitutively active EGFR or EGFR RNAi, in comparison to control. (G) Quantification of junctional (left) and medial (right) Myo-II:mCherry intensity, as shown in F. Statistical significance determined by wilcox-n test, p < 0.05 = *; n = 6 embryos per condition. (H) Representative unwrapped projections showing patterns of anti-β-Catenin (top) and F-Actin (bottom) staining in embryos injected with either DMSO (black) or Cytochalsin D (red). (I)Quantification of junctional F-Actin and β-Catenin intensity, as shown in H. N=3 embryos. Line represents mean plus or minus standard error.

Because actomyosin can also regulate AJ levels and dynamics, we next investigated whether the EGFR-dependent AJ level elevation depended on actomyosin. Cytochalasin D injection in early stage 5 embryos strongly depleted F-actin throughout the embryo, allowing us to assess the effect on AJs (Fig. 6H). We found that AJs still accumulated in a double-sided gradient, even in the absence of junctional F-actin, although it is likely that the structure, function, and dynamics of these junctions is compromised (Fig. 6H – I). To further test whether junctional actomyosin impacts AJ levels, we depleted embryos of Cysts/Wireless/Dp114RhoGEF, an ectoderm-specific RhoGEF that is essential for junctional myosin enrichment in the germband^74,75^. Cysts RNAi did not affect impact the DV AJ gradient, consistent with AJ enrichment not depending on actomyosin (Supplemental Fig. 5A, B). We overexpressed E-Cadherin to determine if altering junction levels regulated junctional actomyosin. Overexpression avoided artifacts that result from loss of tissue integrity when AJs are compromised. E-Cadherin overexpression resulted in elevated actomyosin at subapical junctions and the spreading of actomyosin along the entire lateral contact between cells (Supplemental Fig. 5C). Overall, our data suggested that EGFR regulates AJ levels along the DV axis, which then influences junctional actomyosin levels.

EGF signaling could independently affect junctional actomyosin recruitment and ventral cell movement, or EGF’s effect on ventral cell movement could be driven by junctional actomyosin recruitment. To test this, we again took advantage of the phenotype of depleting the ectoderm-specific RhoGEF, Cysts/Wireless/Dp114RhoGEF. Cysts-RNAi abolished junctional myosin recruitment, as previously described (Supplemental Fig. 5D, compare to Fig. 6F)^74,75^. Cell tracking analysis on Cysts-depleted embryos showed a decrease in ventral cell displacement rate to a similar extent as we observed for EGFR RNAi (Supplemental Fig. 5E, F), compare with Fig. 4E, F). Taken together, these results suggested that EGFR-dependent junctional actomyosin recruitment drives ventral neuroectoderm cell movement towards the ventral midline during the final phase of mesoderm internalization.

## Discussion

The AP and DV patterning systems in *Drosophila* are conventionally thought to mediate distinct morphogenetic processes, germband extension and mesoderm invagination, respectively. Recent work has highlighted possible crosstalk between AP and DV patterning systems. Planar cell polarized junctional myosin, was shown to be present in the mesoderm^41^. In addition, it was shown that AP patterned, pair-rule genes, exhibit variable expression levels along the DV axis, in a manner that matches the DV gradient of myosin^60,76^. However, evidence for a DV patterned gene that clearly regulated convergent extension and mesoderm invagination was lacking. Here, we showed that DV patterned EGF signaling modulates the levels of neuroectodermal junctional actomyosin between AP cell neighbors and mediates the first phase of germband extension and the final phase of mesoderm invagination, suggesting that these morphogenetic movements, while happening in separate domains and under control of different patterning programs, are functionally coordinated. We showed the non-autonomous role for the ectoderm in mesoderm invagination was supported by demonstrating that: 1) EGFR expression and activation was restricted to the ectoderm, 2) EGFR depletion disrupted mesoderm internalization, 3) EGFR activation regulates junctional actomyosin through apical AJ levels, and 4) independently inhibiting the ectoderm specific RhoGEF, Cysts/Dp114RhoGEF, displayed a similar disruption in neuroectoderm ventral displacement as EGFR depletion.

### Intersection of AP and DV patterning regulates Drosophila gastrulation movements

Advances in light-sheet imaging and computational analysis have revealed the choreographed tissue movements that accompany *Drosophila* mesoderm invagination^47,48,59–61,76^. For instance, shortening and stretching of dorsal cells and mitotic rounding have been shown to be responsible for making up much of the surface area lost during mesoderm internalization^47,48^. Ventral movement of the neuroectoderm has been associated with the final stages of mesoderm internalization. This movement of neuroectoderm/germband is associated with the first ‘fast’ phase of germband extension and can at least partially happen in the absence of pulling forces from the mesoderm^47^.

The functional contribution of this ventral movement of the neuroectoderm to mesoderm internalization was poorly understood because mechanisms responsible were unknown. Here, we showed that disrupting AJs and junctional actomyosin in the neuroectoderm through inhibition of the EGFR pathway perturbs completion of mesoderm internalization, demonstrating that these distinct morphogenetic movements are coupled. Consistent with the importance of the connection between mesoderm and ectoderm in the final fast phase of internalization, prior work showed that laser ablations at the boundary between mesoderm and ectoderm, stalled mesoderm internalization^77^. Interestingly, it appears that ventral movement of neuroectoderm during mesoderm internalization requires junctional actomyosin cables enriched between AP neighbors, suggesting that anisotropic pulling forces and possibly multicellular cables promote ventrally-directed neuroectoderm convergence and closure at the ventral midline^60,78–80^.

How is EGFR encoding the modulation of junctional actomyosin activity? Junctional actomyosin is enriched in a ventral-to-dorsal gradient and exhibits planar cell polarity, which predicts tissue flows during gastrulation^26,27,60,78^. Toll-like receptors, Leucine-Rich Receptors, and the adhesion GPCR, Cirl, are all expressed in stripes and are membrane anchored providing a mechanism to promote planar cell polarity by activating signaling at specific cellular interfaces^31–33^. Because germband extension ultimately finishes in the absence of EGFR, it is likely that these other signaling systems are sufficient for PCP and axis extension. Interestingly, EGFR and *rho* gene expression are modulated along the AP axis, exhibiting an AP-striped expression (Supplemental Fig. 1)^67,81^. Given that EGF ligand is released and signals outside its expression domain, we do not favor a model where EGFR signaling cues planar cell polarity. Because myosin enrichment on DV cables depends on mechanosensitive actomyosin recruitment and/or stability and this mechanosensitivity exhibits a DV pattern across the tissue^61,80^, we propose that EGFR is responsible for the patterned mechanosensitive actomyosin recruitment in the germband. Interestingly, the mechanosensitive myosin recruitment does not depend on junction orientation – stretching junctions in AP also recruits myosin, consistent with a soluble cue^61^. Thus, we propose that mesoderm invagination pulls the neighboring neuroectoderm, which due to EGF activity and the mechanical cue, results in myosin recruitment to stretching DV edges, which jumpstarts phase 1 of axis extension while enabling the completion of mesoderm invagination.

### AJs control embryo-wide actomyosin distribution

Our data support a model where EGFR activity regulates junctional actomyosin through the regulation of AJ levels. A previous study showed the actomyosin can cluster AJs leading to their endocytosis^82^. However, our work shows that at the embryo level there is a clear correspondence between AJ and junctional actomyosin levels. Consistent with this, AJs exhibit mechanosensitive enrichment upon actomyosin contractility^14,22^. While we detected a clear role for AJ levels regulating actomyosin, disruption of actomyosin did not eliminate the DV pattern of AJ by either Cytochalasin D injection or Dp114RhoGEF depletion, suggesting that this pattern is initiated and/or maintained independently of actomyosin regulation.

At a cell level, actomyosin, and AJ proteins are planar cell polarized and enriched on complementary interfaces in germband cells^26,78^. Despite being planar cell polarized, AJ proteins are present around the entire circumference of the cell. Therefore, it is possible that AJs along DV interfaces recruit actomyosin. Alternatively, because tricellular junctions between horizontal and vertical interfaces are sites of force transmission required to link multicellular actomyosin cables together^20,80^, it is possible that higher AJs on AP interfaces could also lead to elevated tension and, thus, actomyosin enrichment.

Prior work has shown that perturbations that elevate ectodermal actomyosin and shift ectodermal AJ polarity to resemble mesodermal cells inhibited mesoderm and invagination, such as mutants in *Bearded* (*Brd*) genes^83,84^. One significant difference between the *Brd* mutants and the effect of EGF signaling is that the *Brd* mutant elevated medioapical contractility^83^, whereas EGFR specifically affects junctional contractility. The critical role of junctional myosin in promoting ventral-directed movement of the neuroectoderm was further supported by the Dp114RhoGEF mutant. Increasing junctional contractility in DV oriented actomyosin cables could facilitate mesoderm invagination by propagating ventral forces to promote large scale movement of the tissue along the entire DV axis as observed by light sheet imaging^47,60^. Regardless of the molecular or cellular mechanism, EGFR signaling results in the co-regulation of AJ and junctional actomyosin levels across the embryo, which in this case promotes mesoderm internalization.

### What downstream of EGFR connects the pathway to AJs

There are many possible paths in which the EGFR pathway could regulate AJs and downstream tissue flow. Receptor tyrosine kinase signaling can have multiple downstream outputs, including the Ras-MAPK and the PI-3 kinase branches^85^. Thus, it is possible that multiple pathways operate in parallel to control multiple cell activities. Recent work showed that Toll-2 regulates actomyosin through regulating PI-3 kinase^86^. Our evidence for EGFR suggests that the effect on AJs is at least partially mediated by Ras, but it is possible that EGFR also contributes to PI-3 kinase activation in parallel to Toll-2. Furthermore, PIP3 promotes basolateral protrusive actin activity and it is possible that EGFR could contribute to tissue flow through regulating these protrusions^30^. However, because the EGFR mutant phenotype is phenocopied by Dp114RhoGEF depletion, we favor a model whereby EGFR is acting through apical, junctional actomyosin.

EGFR could also have a more direct regulation of AJs. EGFR has been shown to co-precipitate with the cadherin-catenin complex in *Drosophila* embryo lysates^87^, and also in mammalian cell culture systems^88^. In cell culture, EGFR directly phosphorylates β- catenin and p120-catenin^88^, which in turn can modulate recruitment of actomyosin regulators downstream of EGF stimulation^89^. In Zebrafish, EGFR modulates junctional levels of E-cadherin by tuning the rate of endocytic turnover, which is necessary for tissue flow during gastrulation^90^. Furthermore, tyrosine phosphorylation of the AJ has been shown to regulate AJ levels in the *Drosophila* embryo^91^. Interestingly, the EGFR phenotype of slowing the initial fast phase of extension and reducing both AJs and actomyosin, is similar to that seen in Src mutants^71^. Together this suggests that EGFR may be acting on AJs by modulating the phosphorylation state of its constituent proteins, either through direct phosphorylation by EGFR, or a downstream kinase, such as Src. Another possibility is that EGFR signaling regulates AJs through the small GTPase Rap1. Rap1 and its effector Canoe are required for AJ assembly during cellularization^92^. Rap1 is activated downstream of EGFR in other systems^93^. Interestingly, Rap1 and Canoe function in AJ assembly operates in early gastrulation, but other cues take over later in development^92,94^. This timing coincides with when we observed the most striking regionalization of AJ staining in the embryo. While dissecting the mechanisms by which EGFR regulates AJs through its web of downstream signaling is beyond the scope of this work, the regulatory logic of connecting regionalized signaling to AJs appears to be a general feature of animal development^95^. We show that EGFR plays a pivotal role in *Drosophila* gastrulation and show that EGFR is in a position to integrate both DV and AP patterning processes to coordinate morphogenetic movements.

## Methods

### Fly stocks and genetics

Stocks used in this study are listed in Supplemental Table 1. To generate RNAi knockdown and UAS over-expression embryos, females containing UAS-driven shRNAs against target genes or UAS-driven transgenes were crossed to males carrying a double-maternal driver line with both sqh::GFP and Gap43:mCherry, or sqh::mCherry and E-cadherin::GFP. Resulting larvae were raised at 25°C and females that maternally expressed shRNAs or transgenes and fluorescent markers were crossed to sibling males, and resulting embryos were used for fixation and staining experiments, or live time-lapse microscopy. Unless otherwise noted, mating cages were maintained at 25°C.

### Embryo fixation and antibody staining

Antibodies and their corresponding dilutions used in this study are listed in Supplemental Table 2. For fixed imaging, Oregon-R was used for wild-type. Embryos were first dechorionated in 50% bleach, and then fixed either by a standard formaldehyde fixation, or heat-methanol fixation depending on antibody compatibility [cite previous lab paper]. Embryos were stained with primary antibodies overnight at 4°C, and then with Alexa Fluor-conjugated secondary antibodies for 4 hours at room temperature. After staining, embryos were sliced with a hypodermic needle blade to create cross-sections and mounted in AquaPolymount (Polysciences, Inc.). Images were aquired on a LSM 710 confocal microscope (Carl Zeiss) with a 40x/1.2 Apochromat water immersion objective (Carl Zeiss). In Fig. 6, endogenous mCherry fluorescence was used to visualize Myo-II.

### HCR in situ hybridization

Short antisense DNA probes were designed programmatically using R based on the design principles of HCR v3.0^96^. ∼20-30 probe pairs were designed for each gene including the coding sequence and portions of the 5’ and 3’ UTRs, depending on gene length. Probe sequences are reported in Supplemental Table 3. Probes were ordered as oligo pools (Integrated DNA Technology) and resuspended to 0.5µM in nuclease-free water, and Alexa-conjugated hairpins were acquired from Molecular Instruments. HCR *in situ* hybridization was carried out according to the manufacturer’s protocol for *Drosophila* embryos^96^, and embryos were subsequently mounted in AquaPolymount. Expression patterns were validated against previously published patterns for all genes^97^.

### Time-lapse Microscopy

Dechorionated embryos were mounted ventral-side-up in slide chambers constructed with “embryo glue” (double-sided tape adhesive resuspended in heptane) using #1.5 coverslips as spacers and a #1 coverslip as a cover. After mounting, the chamber was filled with Halocarbon 27 oil (Genesee Scientific). Imaging was carried out at room temperature on a confocal microscope with a 40x/1.2 Apochromat water objective with a pinhole setting between 1 and 2 Airy units using 488-nm argon ion and 561-nm diode lasers. All images were acquired using Zen software (Carl Zeiss).

### Embryo injections

Embryos were first dechorionated and mounted with embryo glue and desiccated for 5 minutes with Dri-Rite (Dririte Company), and covered with a 3:1 mixture of halocarbon 700 and 27 oils. Embryos were then injected with Gefitinib (2.5mg/mL) or cytochalasin D (0.25mg/mL) in DMSO using a microinjection system (World Precision Instruments). Injection of DMSO alone served as a control.

### Image processing and quantitative analysis

Images were processed in FIJI^98^, and data analysis was conducted in R (RStudio). For quantification of junctional enrichment ratios from fixed cross-sections, cross-sections were first “unwrapped” using a polar projection. Intensity was then measured at junctional and lateral positions using line scans of equivalent area, and these values were then divided to produce a trace of normalized junctional intensity. To isolate points corresponding to junctions, the junctional trace was smoothed using a loess function and junctional peaks were then detected using a local maximum filter.

For segmentation of cell membranes or junctions from time lapse movies, data were first pre-processed using histogram-matching bleach correction and a gaussian blur with a radius of 1 pixel. Surface projections were performed using LocalZProjector^99^ and cell segmentation was done using EPySeg^100^. Following segmentation, cell tracking was performed using TissueAnalyzer^101^.

For quantification of junctional versus medial Myo-II enrichment, junctional area was defined based on cell segmentations using E-cadherin:GFP signal, dilated by 2 pixels. Medial measurements were taken using an inverse selection mask (the internal hole within the junctional mask for each cell). Intensity measurements were normalized by subtracting the background cytoplasmic value 5 microns below the apical area.

## Supporting information

Supp. Table 3 - HCR probes

## Acknowledgements

We thank the Developmental Species Hybridoma Bank and Shigeo Hayashi for sharing of antibodies used in this study, and the Bloomington Drosophila Stock Center for providing fly lines. This work was supported by a National Institutes of Health NRSA F32 postdoctoral fellowship to DNC (GM134577), and an NIH MRSA R35 grant to ACM (GM144115). We thank W.J. Nelson and R. O. Hynes and current and former members of the Martin lab for helpful advice and discussion of this work.

## Supplemental Materials

**Supplemental Experimental Procedures:**

**Supplemental Figure 1.**
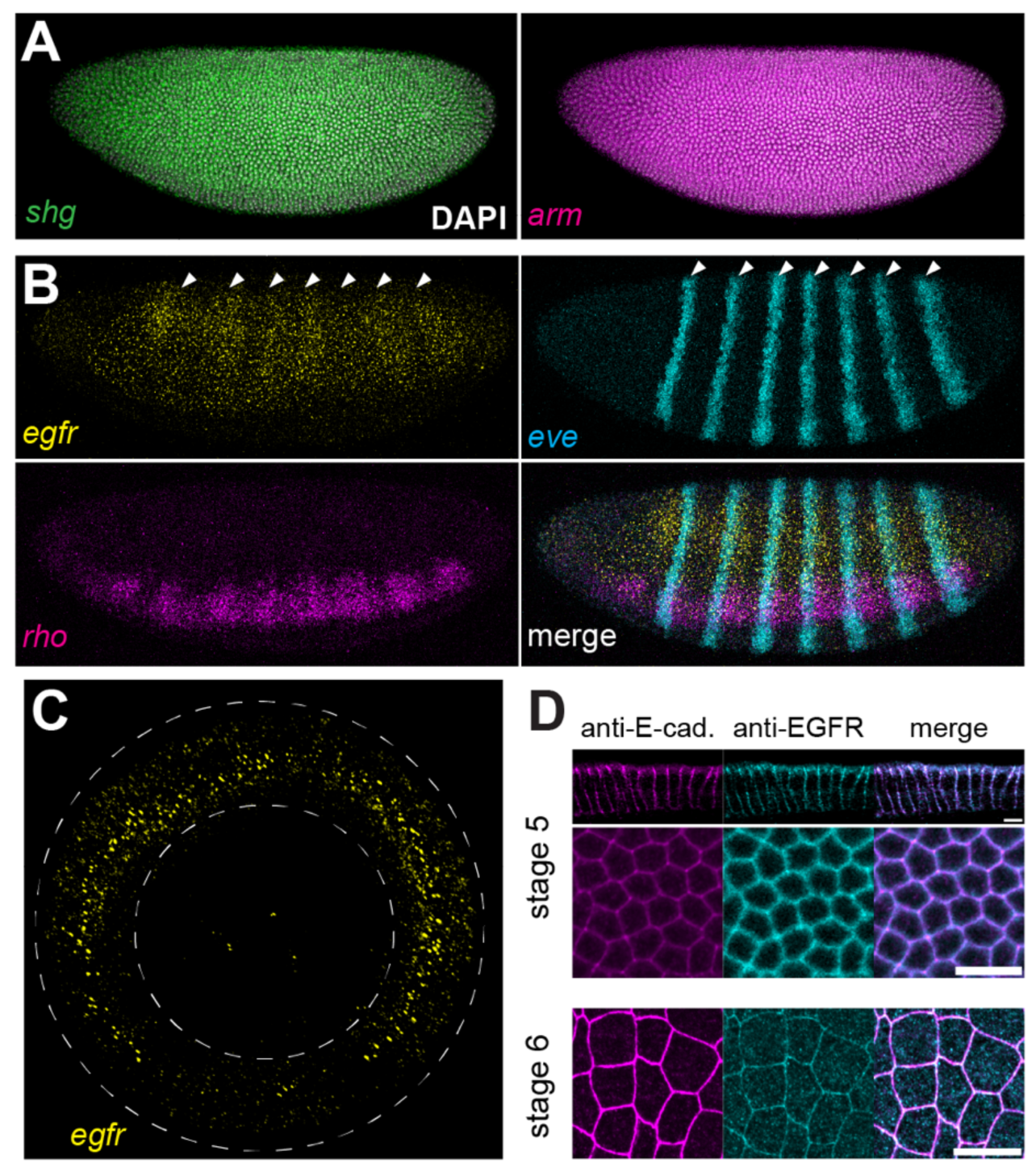
EGFR is expressed in the ectoderm and localizes to AJs and the lateral plasma membrane. (A) HCR *in situ* hybridizations for *e-cadherin* (*shotgun; shg*) and *β-catenin* (*armadillo; arm*), showing ubiquitous expression throughout the embryo at stage 5 (zygotic expression). (B) HCR *in situ* hybridizations for *egfr*, *even-skipped* (*eve*), and *rhomboid* (*rho*) at stage 5 (zygotic expression). Pair-rule stripe pattern is indicated with white arrowheads. (C) Cross-sectional view of *egfr* HCR, showing lower signal in the mesoderm. (D)Antibody staining for E-Cadherin and EGFR in stage 5 and 6 embryos. Scale bar is 10 microns.

**Supplemental Figure 2.**
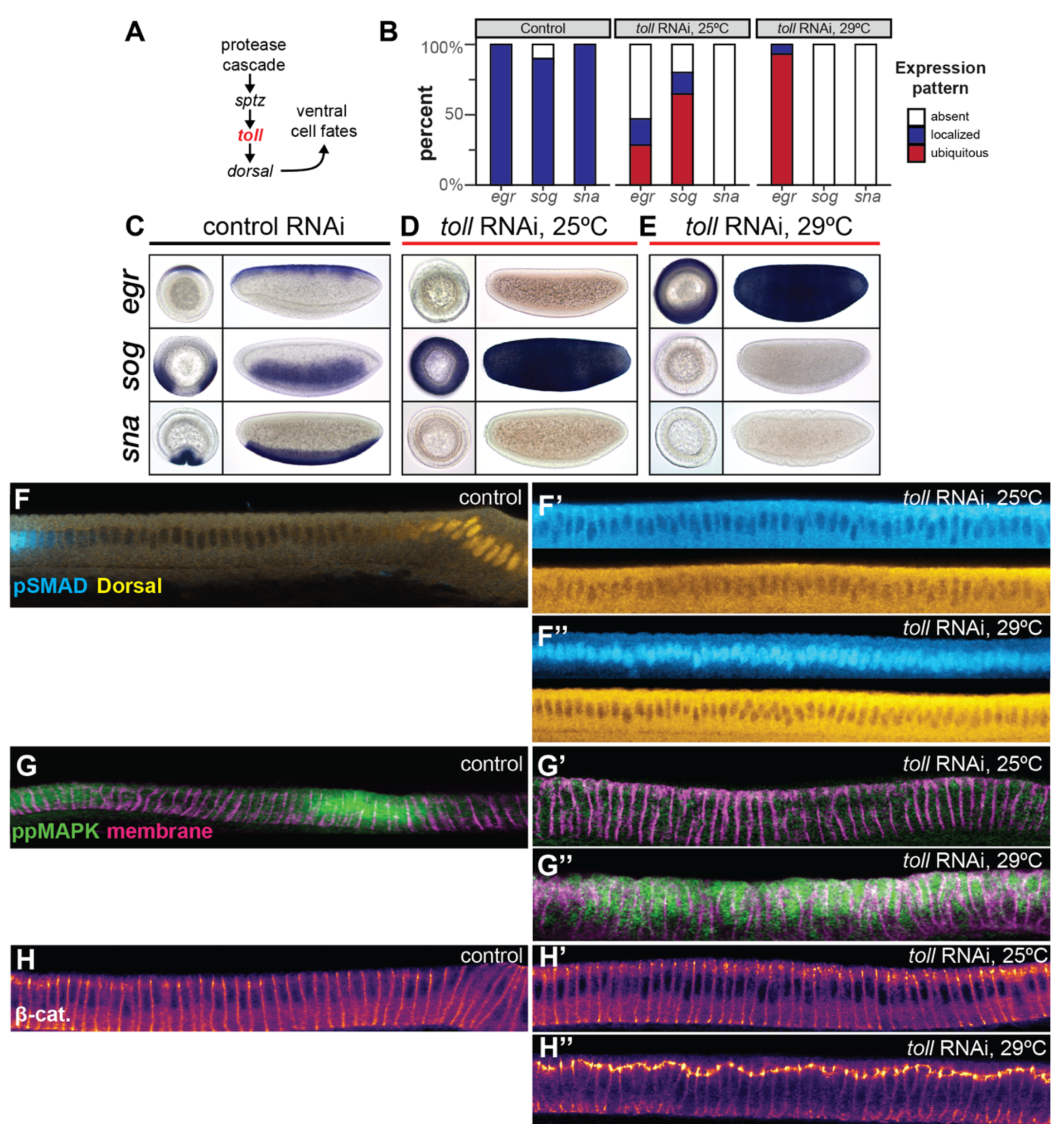
Levels of AJ proteins are downstream of the dorsal-ventral patterning pathway. (A) Schematic of Toll DV signaling pathway. (B)Quantification of gene expression pattern frequencies in lateralized (*toll* RNAi 25°C) or dorsalized (*toll* RNAi 29°C) embryos in comparison to controls. n = 25 embryos per stain, per condition. (C-E) Representative chromogenic *in situ* hybridizations for markers of dorsal (egr), lateral (sog), and ventral (sna) DV expression domains under control, lateralized, and dorsalized conditions. (F)pSMAD and Dorsal antibody staining showing dorsal and ventral cell identities, respectively in control (F) in comparison to lateralized (F’) and dorsalized (F’’) conditions. (G) Di-phospho MAPK staining in lateralized (G’) and dorsalized (G’’) conditions. (H) Anti-β-Catenin staining in lateralized (H’) and dorsalized (H’’) conditions.

**Supplemental Figure 3.**
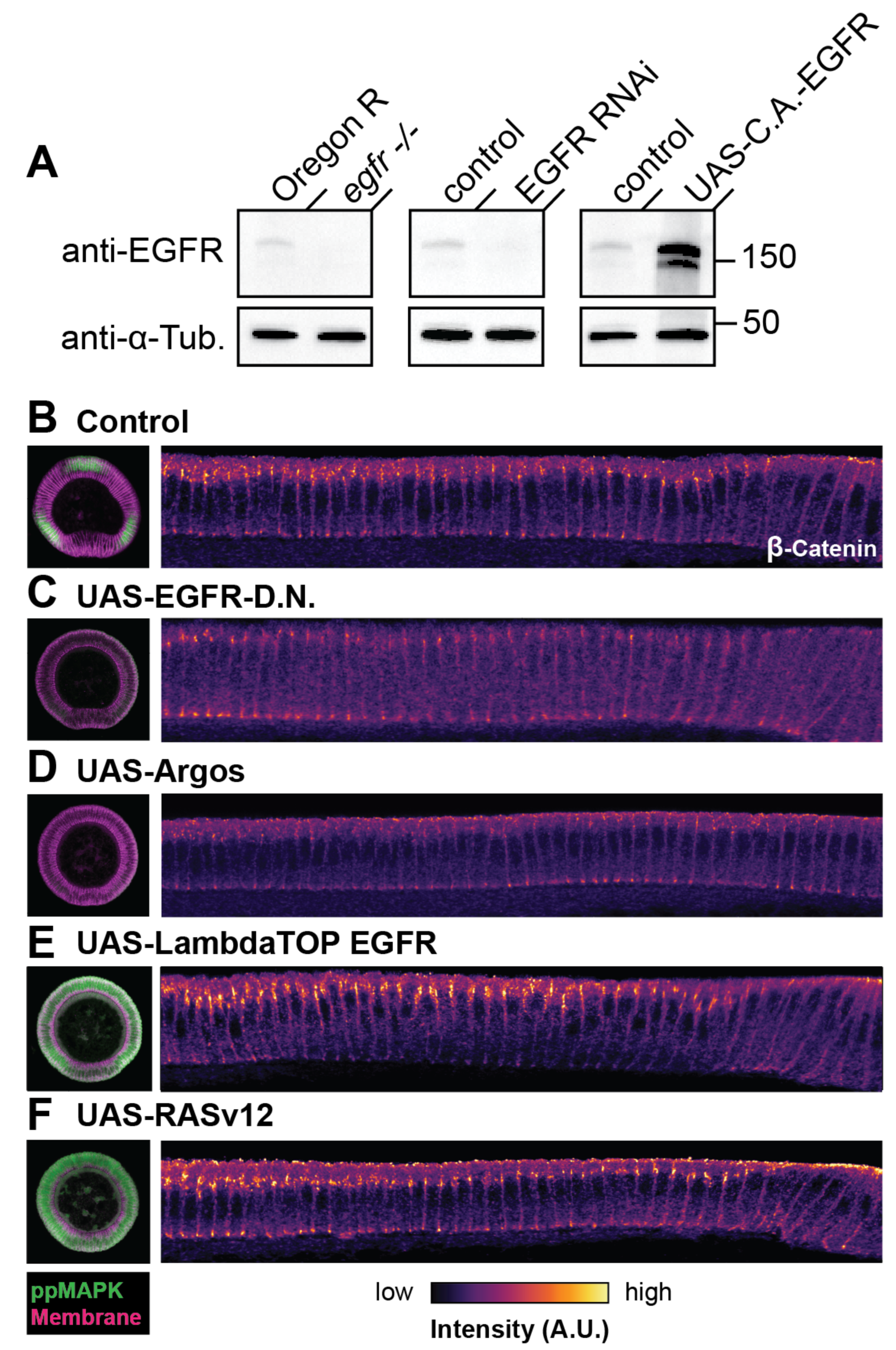
Western blot of EGFR perturbations and additional perturbations that modulate AJ levels. (A) Western blot showing abundance of EGFR protein in 2-4 hour embryos in wildtype, EGFR zygotic null mutant, EGFR RNAi, and EGFR over-expression conditions. Anti-tubulin is shown as a loading control. (B-F) Representative unwrapped projections for control (B) versus additional EGF-inhibiting (C,D) and EGF-activating (E,F) conditions stained for Anti-β-Catenin.

**Supplemental Figure 4.**
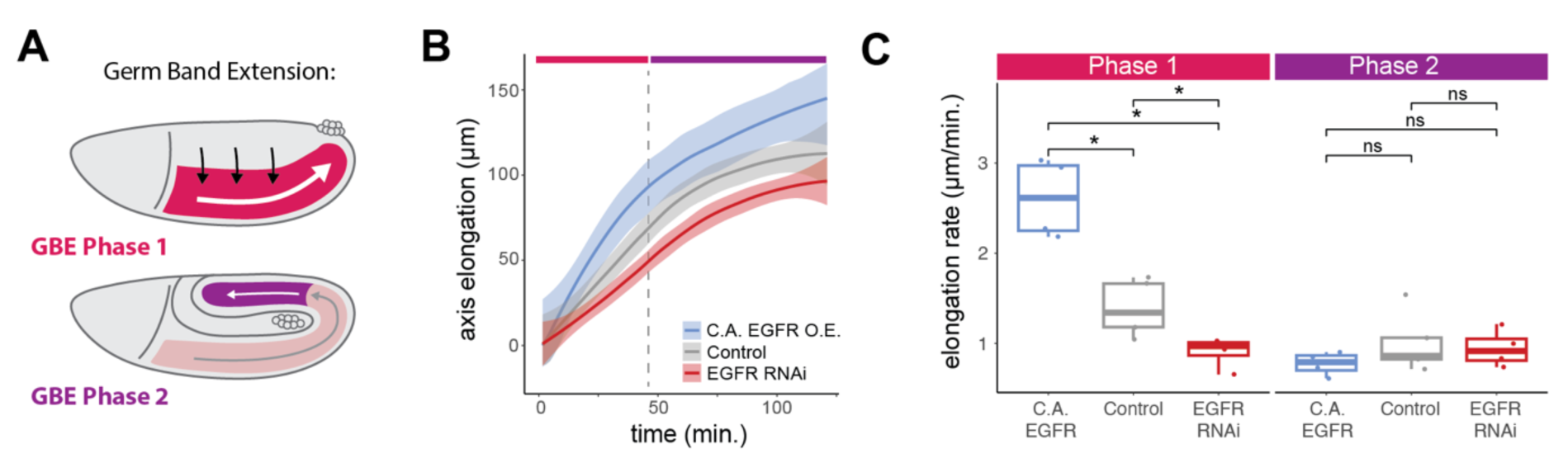
EGFR perturbations modulate the rate of Phase I of germ band extension. (A) Schematic of the germ band extension (GBE) process, showing Phase I (fast) and Phase II (slow). Arrows indicate approximate direction and speed of cell movement. (B) Quantification of GBE, as measured as axis elongation over time, from brightfield microscopy movies in embryos over-expressing constitutively active EGFR (blue) or EGFR RNAi knockdown (red), in comparison to control embryos (gray). Bars and dashed line indicate GBE phases and approximate time of transition, respectively. N=4 embryos per condition. Line represents mean plus or minus standard deviation. (C) Comparison of GBE rates as measured in B, divided into Phase I (left) and Phase II (right). Statistical significance determined by a wilcox-n test, p < 0.05 = *; n = 4 embryos per condition.

**Supplemental Figure 5.**
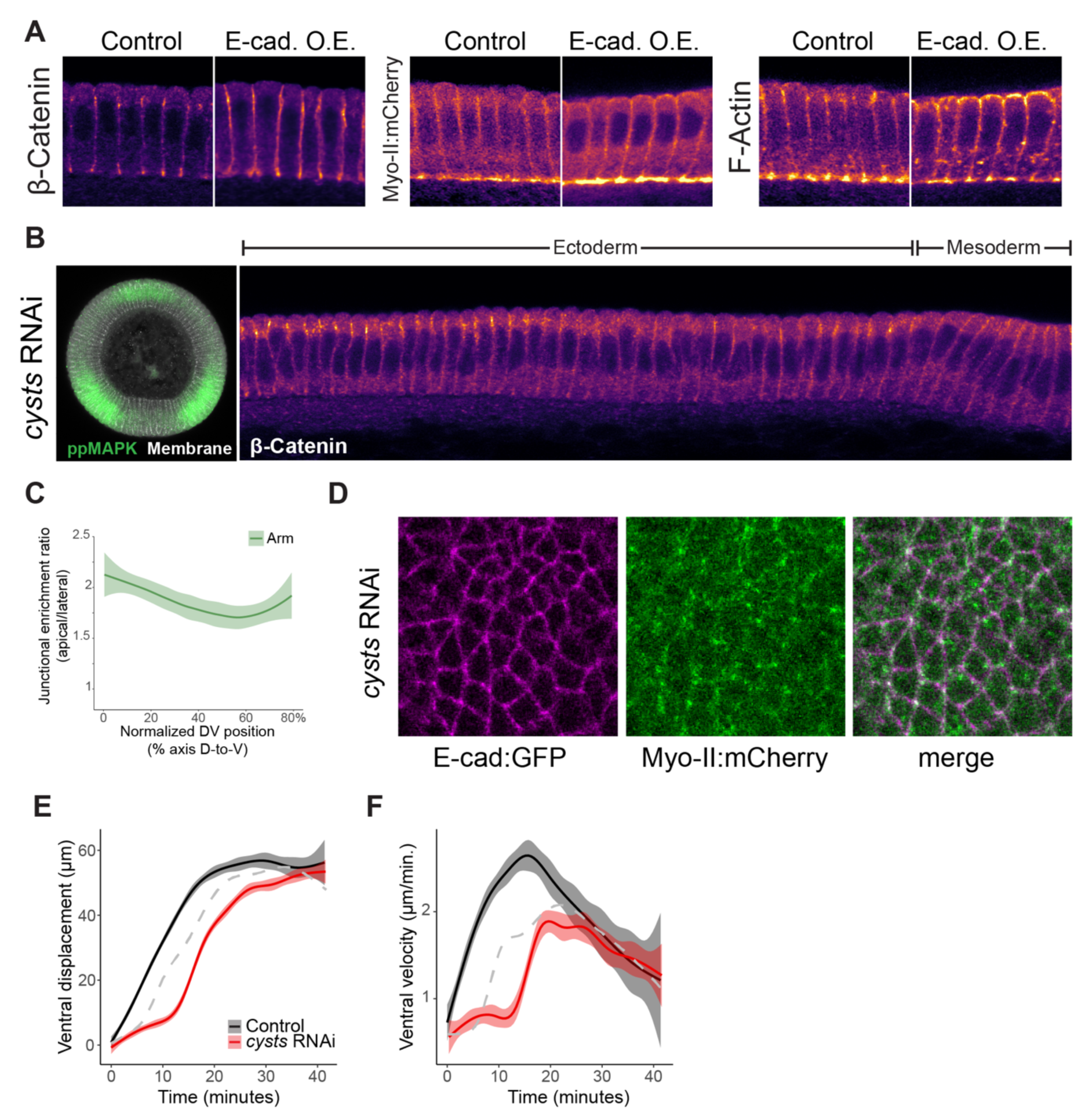
Supplementary to Figure 6. (A) Over-expression of E-Cadherin drives ectopic localization of actomyosin. (b) Representative unwrapped projection of a Cysts RNAi embryo stained for Anti-β-Catenin. (C) Quantification of junctional β-Catenin intensity in Cysts RNAi embryos. N = 5 embryos. Line reperesents mean plus or minus standard error. (D) Representative *en face* view of E-Cad:GFP and Myo-II:mCherry signal in the germ band of *cysts* RNAi during convergent extension. (E-F) Quantification of ventral displacement (E) and velocity (F) over time in ventrolateral aspect movies of *cysts* RNAi embryos (red) in comparison to control (black). Dashed grey line indicates EGFR RNAi trendline from Figure 4E-H. N = 3 embryos; lines represent mean plus or minus standard deviation.

**Supplemental Table 1.**
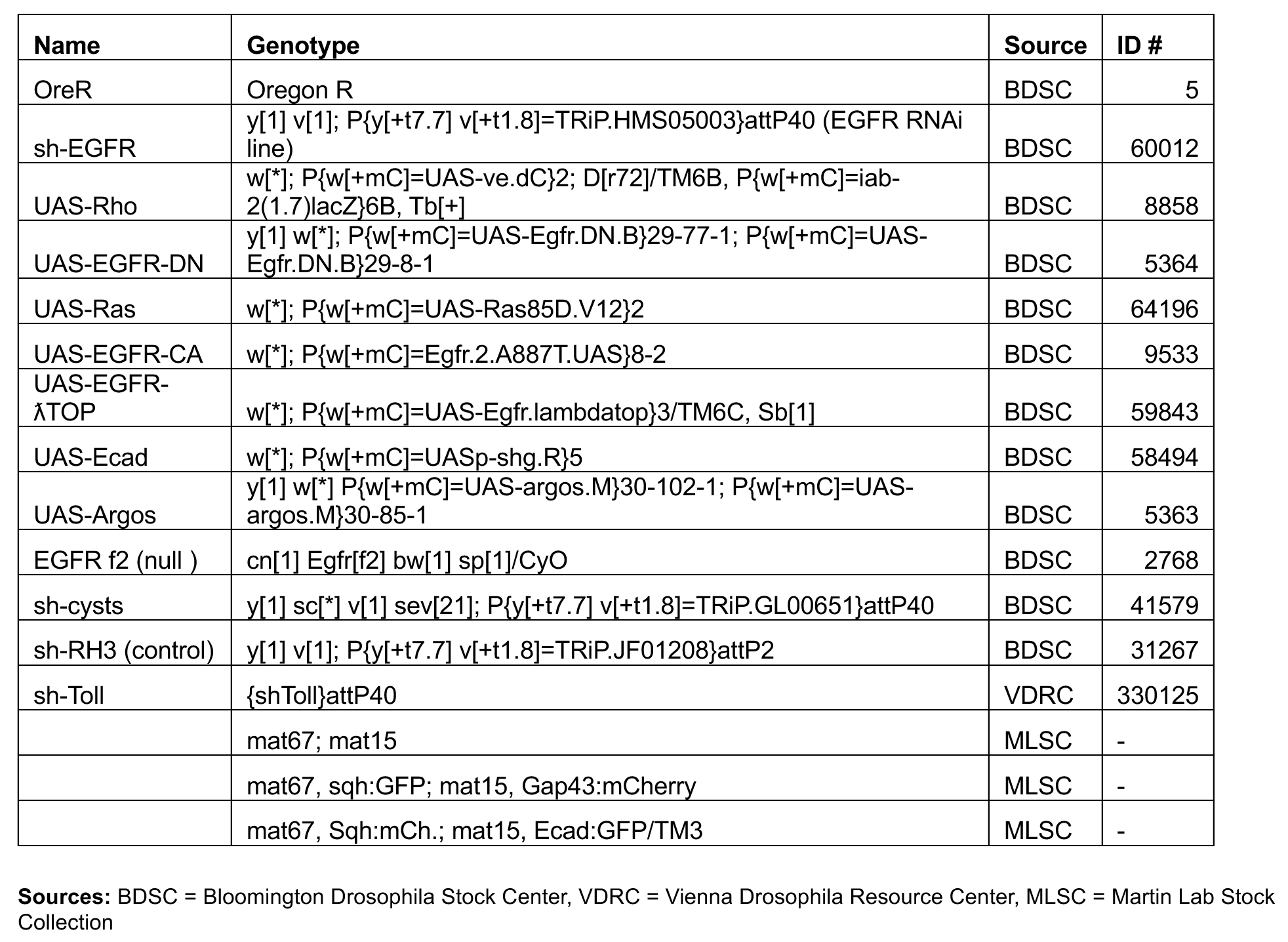
Drosophila lines used in this study.

| Name | Genotype | Source | ID # |
| --- | --- | --- | --- |
| OreR | Oregon R | BDSC | 5 |
| sh-EGFR | y[1] v[1]; P{y[+t7.7] v[+t1.8]=TRiP.HMS05003}attP40 (EGFR RNAi line) | BDSC | 60012 |
| UAS-Rho | w[*]; P{w[+mC]=UAS-ve.dC}2; D[r72]/TM6B, P{w[+mC]=iab-2(1.7)lacZ}6B, Tb[+] | BDSC | 8858 |
| UAS-EGFR-DN | y[1] w[*]; P{w[+mC]=UAS-Egfr.DN.B}29-77-1; P{w[+mC]=UAS-Egfr.DN.B}29-8-1 | BDSC | 5364 |
| UAS-Ras | w[*]; P{w[+mC]=UAS-Ras85D.V12}2 | BDSC | 64196 |
| UAS-EGFR-CA | w[*]; P{w[+mC]=Egfr.2.A887T.UAS}8-2 | BDSC | 9533 |
| UAS-EGFR-λTOP | w[*]; P{w[+mC]=UAS-Egfr.lambdatop}3/TM6C, Sb[1] | BDSC | 59843 |
| UAS-Ecad | w[*]; P{w[+mC]=UASp-shg.R}5 | BDSC | 58494 |
| UAS-Argos | y[1] w[*] P{w[+mC]=UAS-argos.M}30-102-1; P{w[+mC]=UAS-argos.M}30-85-1 | BDSC | 5363 |
| EGFR f2 (null ) | cn[1] Egfr[f2] bw[1] sp[1]/CyO | BDSC | 2768 |
| sh-cysts | y[1] sc[*] v[1] sev[21]; P{y[+t7.7] v[+t1.8]=TRiP.GL00651}attP40 | BDSC | 41579 |
| sh-RH3 (control) | y[1] v[1]; P{y[+t7.7] v[+t1.8]=TRiP.JF01208}attP2 | BDSC | 31267 |
| sh-Toll | {shToll}attP40 | VDRC | 330125 |
|  | mat67; mat15 | MLSC | - |
|  | mat67, sqh:GFP; mat15, Gap43:mCherry | MLSC | - |
|  | mat67, Sqh:mCh.; mat15, Ecad:GFP/TM3 | MLSC | - |

**Sources:** BDSC = Bloomington Drosophila Stock Center, VDRC = Vienna Drosophila Resource Center, MLSC = Martin Lab Stock Collection

**Supplemental Table 2.** antibodies used in this study:

| Antibody | Species | Fixation | Dilution | Source |
| --- | --- | --- | --- | --- |
| anti-E-cadherin | Rt | FAM | 1:50 | DSHB (#DCAD2) |
| anti-Armadillo | Ms | HM | 1:500 | DSHB (#N27A1) |
| anti-Alpha-cat. | Rt | HM, FAM | 1:50 | DSHB (#DCAT1) |
| anti-P120-cat. | Ms | FAM | 1:50 | DSHB (#P1B2) |
| anti-EGFR | Ms | FAM | 1:100 | Sigma (#E2906) |
| Anti-Neurotactin | Ms | HM, FAM | 1:100 | DSHB (#BP106) |
| Anti-Rho | Gp | FAM | 1:500 | Kind gift of Shigeo Hayashi |

**1° antibody species:** Ms = mouse, Rb = rabbit, Rt = rat, Gp = Guinea pig
**Fixation:** HM = heat-methanol fixation, FAM = formaldehyde fixation, MeOH devitellization, FA = formaldehyde fix, manual devitellization
**Sources:** DSHB = Developmental Studies Hybridoma Bank, University of Iowa

